# Wild lab: A naturalistic free viewing experiment reveals previously unknown EEG signatures of face processing

**DOI:** 10.1101/2021.07.02.450779

**Authors:** Anna L. Gert, Benedikt V. Ehinger, Silja Timm, Tim C. Kietzmann, Peter König

## Abstract

Neural mechanisms of face perception are predominantly studied in well-controlled experimental settings that involve random stimulus sequences and fixed eye positions. While powerful, the employed paradigms are far from what constitutes natural vision. Here, we demonstrate the feasibility of ecologically more valid experimental paradigms using natural viewing behavior, by combining a free viewing paradigm on natural scenes, free of photographer bias, with advanced data processing techniques that correct for overlap effects and co-varying nonlinear dependencies of multiple eye movement parameters. We validate this approach by replicating classic N170 effects in neural responses, triggered by fixation onsets (fERPs). Importantly, our more natural stimulus paradigm yielded smaller variability between subjects than the classic setup. Moving beyond classic temporal and spatial effect locations, our experiment furthermore revealed previously unknown signatures of face processing. This includes modulation of early fERP components, as well as category-specific adaptation effects across subsequent fixations that emerge even before fixation onset.

## Introduction

The EEG correlates of face processing have been studied widely over the last decades, as faces represent an important stimulus category in our everyday life. Using well-controlled experimental paradigms, numerous studies have revealed a face-specific modulation of event-related potentials (ERPs) that occur in occipito-temporal electrodes around 170ms after stimulus onset (N170) [1]. Most studies [2–4] (but also see [5]) report this component to be more negative for trials that presented a face, in comparison to other categories like cars, butterflies, or clocks [3].

While the N170 is a highly robust experimental finding, most of what we know about the neural correlates of face processing is derived from ‘classic’ experimental paradigms derived to enable maximal control over stimulus parameters. These include stimulus’ contrast [2], spatial frequency [6], inversion [7], shape [8], integrity [9], or orientation [10]. While more natural stimulus material with varying perspectives and backgrounds [11,12] or movement [13] have successfully been used to produce face-related EEG responses, most experimental setups remain highly artificial. For example, they rely on randomized sequences of stimulus presentations and do not allow for eye movements, although the latter play a central role in natural vision.

A potential consequence that comes into play in free viewing studies are sequential effects. Previously, these effects have been well described, for instance, in choice biases in behavior [14-16], pupil dilation[17], or face identity perception [18]. However, there are also direct effects, i.e. autocorrelations, within sequences of eye movements. One prominent example is the overabundance of forward saccades [19]. Effects of such serial depenfdencies on neuronal activity have been found to occur in early visual areas [20] and higher cortical areas. In an intracranial EEG study, Körner et al. [21] even showed sequential effects for fixation locked ERPs in a visual search task. We will analyze the sequential effect of fixation history on face processing by explicitly modeling the fixation history at the previous fixation locations.

Here, we advance the study of face perception by introducing an experimental and analysis paradigm that allows for active vision on natural scenes. This is accomplished by a combination of three elements. First, we perform simultaneous recordings of eye movements and electrophysiological data. Second, we use an unrestricted free viewing paradigm on natural stimuli, sampled without photographer bias from an HD head-cam that volunteers wore while moving in the real world. Third, we employ a novel analysis pipeline of fixation-ERPs that is capable of controlling for temporal overlap in neural processes elicited by rapidly occurring eye movements as well as disentangling and adjusting for the effects of varying eye movement parameters.

Previewing our results, we demonstrate a high within-subject correlation of N170 effect sizes across free viewing and a classic experimental paradigm, validating our approach. Importantly, we observe a reduction in the effect size and its variance across subjects for free viewing, indicating that the more natural setup led to more consistent brain activity. In addition to these N170 effects, the free viewing condition shows a face-selective modulation already at the P100.

Moreover, we also find evidence for sequential effects in subsequent fERPs, emerging even before fixation onset. These findings highlight the importance of understanding eye movements as a sequence of peripheral preview and foveated analysis and not as a series of independent, rapid stimulus onsets, and add further support for utilizing more natural stimulus paradigms.

## Results

To better understand the benefits and limitations of both classic laboratory and more natural experimental setups, we recorded high-density EEG from a group of participants in two main conditions: in the classic lab condition, we conducted a traditional lab study of face processing, showing faces and objects while participants maintained central fixation. In the free viewing condition, we allowed for free eye movements and used natural stimuli as sampled from an HD head-cam that volunteers wore while moving in the real world.

### The free viewing paradigm replicates classic face processing ERPs, while reducing the cross-participant variance

The classic lab condition contrasts photographs of isolated faces and objects. We observe well-known signatures of face processing. These are predominantly visible as a negative ERP peak at around 170ms (Fig 1A), with face trials showing a more negative deflection than trials on objects at individual N170 peaks (Faces: mean individual maximum of −3.5μV 95%-CI: [-4.6;-2.3μV], Objects: −0.8μV [-2.1;0.3], difference: −2.7μV [-3.5;-2.1]; further comparisons in supplementary Table 1). We do not see a difference in the P100 peak amplitudes (Faces: 6.0μV 95%-CI: [4.7;7.3], Objects: 6.2μV [4.9;7.6], difference: −0.2μV [-0.5;0.1]). These results qualitatively agree with early reports of the N170, and are quantitatively even more pronounced [7].

**Fig 1.**
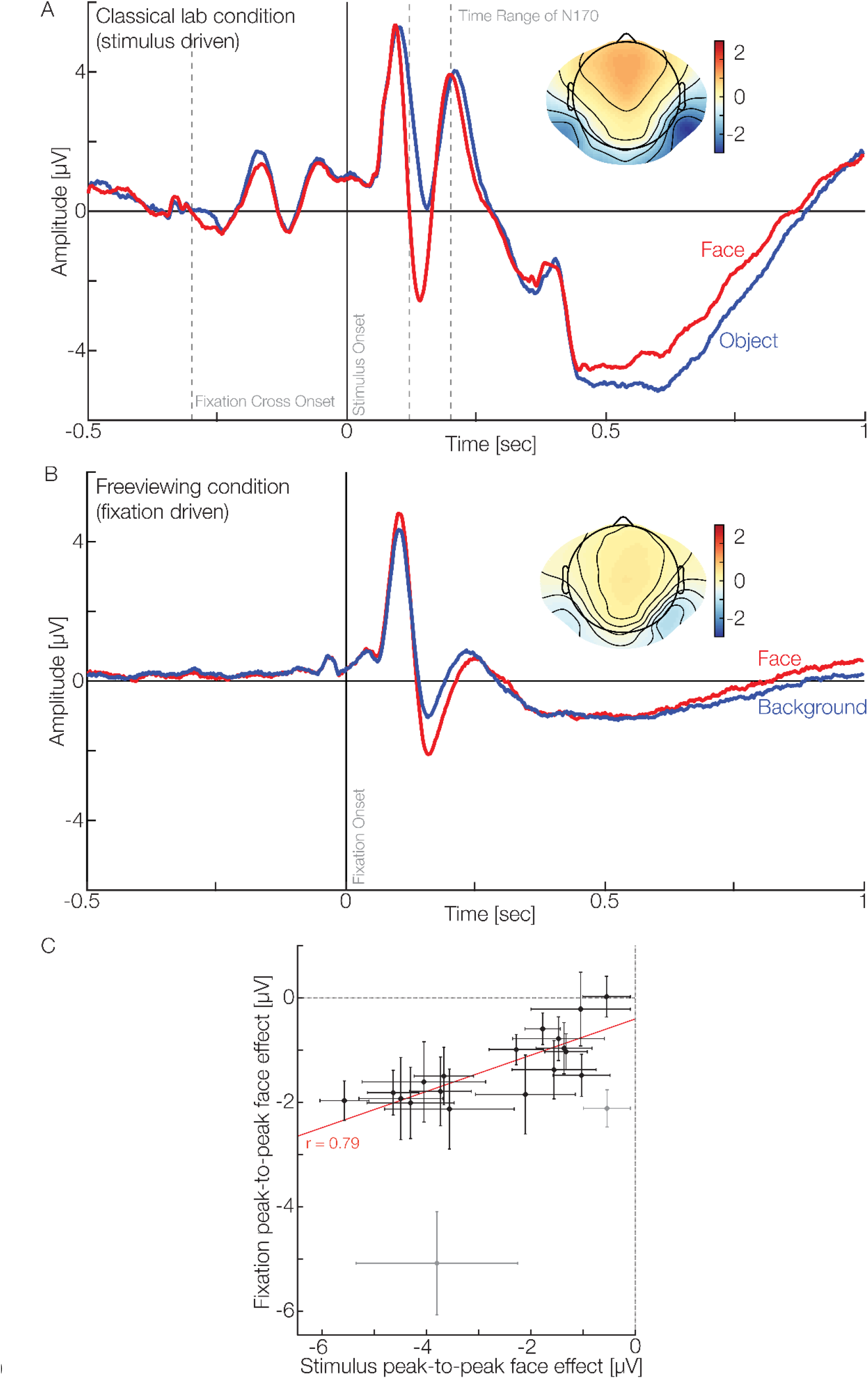
ERPs of the passive and free viewing condition and their correlation displayed with average reference. A. Stimulus-driven ERP of the passive condition. With our experiment, we can reproduce previous findings of the N170 being larger when faces are presented. The topographic plot visualizes the average activity of the N170 time range for all electrodes. B. Fixation-driven fERP of the free viewing condition. Here we can see that fixations on a face produce a more negative N170 than those on the background. Additionally, the topography shows a generally weaker activation but the same parieto-occipital pattern of stronger right lateralization. C. Correlation of the peak-to-peak effects. The peak-to-peak differences (amplitudes of P100 -N170, face trials - non-face trials) in the passive and active conditions correlate (r=0.79). Grey data points were automatically excluded by the robust statistics toolbox. Please note that all data shown in this plot are corrected for overlap and eye-movement-dependent effects.

Next, we investigate fixation-related ERPs in the free viewing condition on natural scenes. During a 6-second trial, participants were allowed to explore scenes photographed in a local shopping mall, while recording eye tracking data and EEG. Each scene contains one to seven faces. This enables us to classify fixations as being on human faces, the scene’s “background” or “other”. Because neural activity from subsequent fixations can overlap in time, we perform a linear deconvolution using the unfold toolbox [22]. To account for systematic differences in conditions between saccade amplitude and the fixations’ horizontal and vertical position, we model several eye movement-related covariates as non-linear effects using spline regressors [22]. Moreover, we model several sequential effects. A full specification of the model can be found in the Methods section.

Collapsing over sequential effects, and controlling for temporal overlap and the effects of eye movements, we observe that the fixation-induced ERP (fERP) is modulated by fixations on faces. In particular, the N170 is more negative for fixations on a face than for those on the background (Fig 1B, Faces: −2.5μV 95%-CI: [-2.9;-2.2], Background: −1.4μV [-1.7;-1.1], difference: −1.1μV [-1.5;-0.9]). At the same time, the more natural paradigm leads to a stronger P100 when a face is fixated (Faces: 5.2μV 95%-CI: [4.3;6.3], Background: 4.8μV [3.9;5.8], difference: 0.4μV [0.3;0.7]; further comparisons in supplementary Table 1). In summary, these findings replicate previous passive presentation experiments in a more natural setting, but also provide evidence that, under such natural viewing conditions, additional effects occur at earlier processing stages.

In addition to the presence of face-related N170 effects in both paradigms, we perform a more stringent test of the statement that the same face-related brain processes are at play and correlate the effect sizes in both conditions across participants (Fig 1C). Indeed, a robust skipped Pearson correlation of the peak-to-peak N170 effect shows a strong correlation (r=0.79 [0.63;0.92]). Assuming a perfect correlation and taking into account the within and between-subject noise estimates (noise ceiling, see Methods), this value is within the upper bound of observable correlations. These results show that participants with a stronger N170 effect in the classic lab condition also show a stronger N170 effect in the more naturalistic free viewing condition, meaning that individual differences generalize to more ecologically valid setups. Interestingly, we do not only observe smaller between-condition differences, but also a lower between-subject variance in the free viewing condition than in the classic lab condition (with a standard deviation of 0.6μV [0.5;0.9] and 1.5μV [1.3;2.0] respectively, difference [0.5;1.2]). Investigating the effect sizes between the experiments leads to non-significant differences (Cohen’s d of Passive Viewing 95%-CI [1.3;2.2], Free Viewing [1.3;3.2], difference [-1.5;0.2]). This result suggests that the absolute size of the variance in more natural settings is more stable across subjects.

### Unrestrained spatiotemporal analyses reveal further effects of face processing across subsequent fixations

Having verified our experimental and analysis approach, we expand our analyses beyond the commonly used, yet restricted set of electrodes and time windows. This allows us to analyze the complete temporal dynamics of face processing across all electrodes. To correct for the massive multiple comparisons problem across time and sensors we use threshold-free cluster enhancement on the single subject parameter estimates resulting from our unfold model (see Methods). A benefit of our free viewing paradigm, compared to experiments requiring participants to fixate, is that sequential effects across voluntary fixations can be investigated. In addition to the object category viewed at the current fixation (face vs. background), we model whether a fixation was previously on a face. Including this sequential predictor in the model further allows us to investigate the interactions between the current and the previous fixation category. Finally, we include the influence of gaze shifts within a single face and between different faces (see Methods/Fig 5B).

Based on our analyses of the model, we observe a significant difference (cluster permutation test with TFCE-corrected *α*=0.05) in the main effect for the current fixation type, face vs background, beyond the commonly investigated effect time and location. This difference is likely driven by two clusters from 74 to 242ms, in frontal parieto-occipital and occipital electrodes (Fig 2A, thick black lines, and thick black circles). Temporally, two components can be distinguished: An early positive P100 at occipital electrodes with a topography implicating processing in the early visual regions (peaking at 1.24μV) and a later bilateral N170 effect, dominant at parieto-occipital electrodes (peaking at −1.65μV). Both source configurations are accompanied by their respective frontal dipole-counterpart, which for the N170 is often termed the VPP. These results support our previous findings and further demonstrate that voluntary fixations on natural faces lead to earlier differences, including the timeframe of the P100.

**Fig 2.**
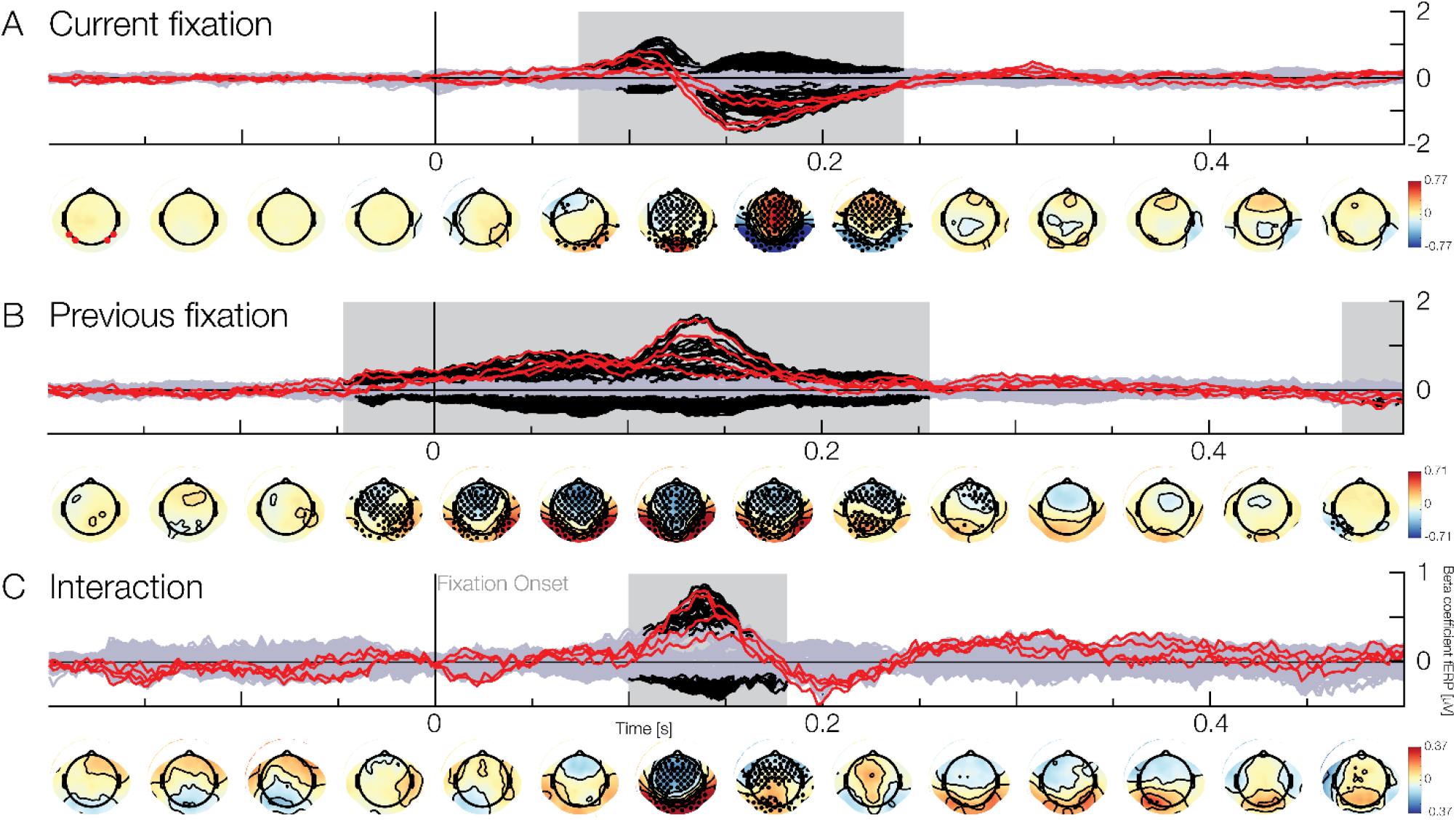
Model results of the free viewing condition (red lines indicating the beta for the classic N170 electrodes). (A) Effect of the current fixation. When currently fixating a face, the amplitude will be stronger in the P100 and N170 range (black lines and black dots). Channels marked in red are P7/8 and PO7/8. (B) Effect of the previous fixation. When saccading from a face, the amplitude will be more positive in parieto-occipital and more negative in fronto-central electrodes. This effect is already present before fixation onset until after the N170. (C) Interaction of the current and previous fixation. When participants saccade between two faces, the ERP will be significantly decreased during the N170. All effects here were modeled using effects coding (−0.5 for background, 0.5 for faces).

In addition to analyses of the main effects of face processing based on the category of the *currently* fixated object, we next analyze the main effect of the *previous* fixation. That is, we investigate the difference between fixations coming from the background versus those coming from a face, regardless of the currently fixated category (the classical ERP plot can be found in supplementary Fig.1). This reveals significant effects that originate from clusters in frontal and parieto-occipital electrodes (Fig 2B). Notably, the cluster starts about 50ms before fixation onset and extends up to 256ms after fixation onset. The cluster topography implies the same source configuration as the N170, but as an inverted effect: more positive in parieto-occipital and more negative in frontal electrodes (peaks at 1.7μV and −0.62μV respectively). Together with the main effect observed for the current fixated category, this means that not only is the N170 less strong when previously fixations were on a face, but that this modulatory effect appears already before the new fixation started. A shorter second cluster shows effects between 469ms and 510ms (peak at −0.43μV) in a small set of electrodes. To conclude, when performing a saccade coming from a face, the EEG activity elicited by the current fixation will be more positive in typical N170 sources, even before the current fixation onset, clearly indicating sequential effects across subsequent fixations.

Having investigated the two main effects, the category of the current and previously fixated objects, we examine them in light of their interaction. Testing the interaction term reveals a significant cluster with positive and negative activations from 100ms to 182ms (Fig 2C). The negative betas are strong in frontal electrodes with a peak of −0.43μV, while the positive betas, peaking with 0.88μV, are located at parieto-occipital electrodes. Combining this with the two main effects of previous and current fixation, this result implies that the N170 has a smaller amplitude (i.e. is more positive) if a participant saccades between two faces. This effect can be understood in terms of neural adaptation effects. Notably, the early part of the main effect of the previous fixation shows no co-occurring interaction. Thus, the early part of the main effect of previously fixating a face seems to resemble a reactivation of the previous fixation type and the potential adaptation effect is therefore limited to the neuronal substrate activated relatively late in the process.

While our previous analyses focus on fixations between the background of a scene and faces, or across separate faces, some fixation sequences appear within the same face (Fig 3). Analyzing these data, we observe a significant interaction including a first cluster located parieto-occipitally, starting in electrodes on the right hemisphere at around −75ms and spreading bilaterally over time until 146ms, peaking at −3.41μV (frontal electrodes peak at 1.88μV). A second weak and short cluster around 475ms is located in the right occipito-temporal electrodes starting at 449ms until 490ms but due to the distance to the fixation event, this remains difficult to interpret. In summary, performing two consecutive fixations on the same face will lead to a weakened ERP signal between the saccade onset and the P100, indicating a preview and adaptation effect.

**Fig 3.**
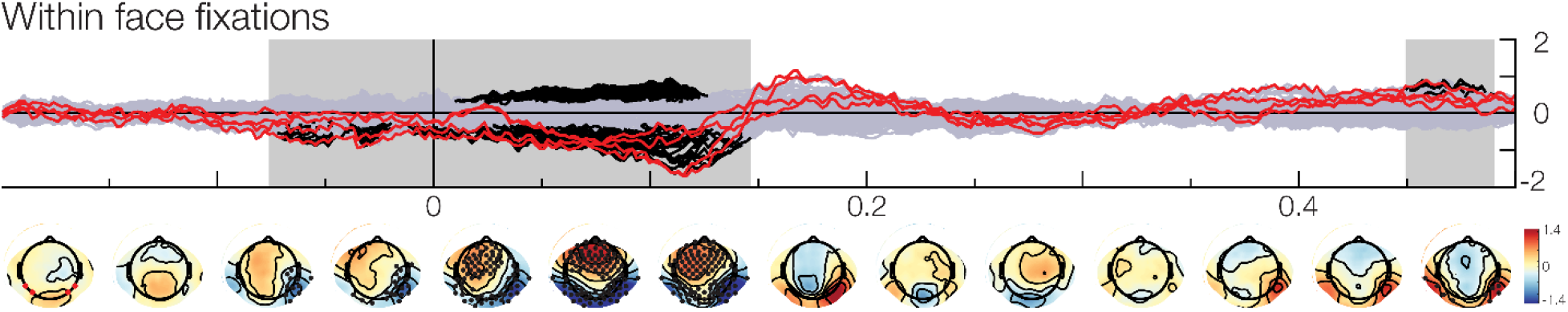
Results of saccading within the same face. When consecutive fixations are made within the same face, the activation will be weaker even before fixation onset starting in electrodes normally associated with the N170. This indicates an adaptation effect up until the N170. Please note that even though the betas show an opposite behavior to Fig 4, the effect is the same, due to the coding in our model. This interaction is coded with 0 for non-faces and 1 for faces when saccading within the same bounding boxes.

## Discussion

### Reproducing and extending the classic observations

The main goal of this study is the validation and extension of classic experimental results under more naturalistic experimental conditions. While previous studies used naturalistic setups, they either employed everyday stimulus material, but lacked eye movements [11,13] or they allowed for eye movements but used artificial stimulus material [24,25]. These studies advanced face perception research, but lacked the crucial combination of embodiment and natural stimulus material. Previous literature showed that the neural correlates of perception differ between passive and active perception [24,26,27] and that naturalistic stimuli will lead to different activation from artificial ones [13,28]. As the combination of these two aspects is what we encounter in our everyday life, it is necessary to combine them to obtain the full picture of naturalistic face perception. Not only are we able to confirm results classically reported in passive perception experiments[3] but we also replicate the findings of one of the first studies combining free viewing and face perception [24] comparing fERPs recorded during task-driven free viewing to ERPs in a passive stimulation task using cutout faces. Yet, we could demonstrate that naturalistic face processing will lead to earlier effects, including time points classically defined as the P100, and extend beyond parieto-occipital electrodes, throughout the whole scalp. This indicates extensive processing including a strong activation of the underlying neuronal sources like the posterior STS and the FFA [29] (see [30] for a review). However, our results contrast those of Soto et al. [25]. In their study, subjects freely viewed a real-world stimulus display containing cutout faces. In addition, Soto et al. did not control for eye movement-related parameters or overlap. Thus, the difference to our study might be due to the difference in defining the fixation onset and their lack of statistical control for eye movement parameters. To conclude, here we replicate the classic findings in more natural settings and further extend these observations to larger time ranges including the early parts of the visual response.

### Similar processes in classic and the more naturalistic conditions

Whether our fixation and more traditional stimulus-evoked responses describe the same processes is an important question. While previous studies have shown that face processing related eye movements generalize from lab-based settings to mobile recordings [31], this generalization is especially debated for the P100 and the lambda response, but also the N170 and the N1 of the lambda complex [31]. Here we focused on the correlates of face processing, as described by the difference in the P100/N170 peak-to-peak amplitude. Still, our study adds to this discussion, as we found a high correlation of 0.78 between eye-movement N1 and traditional N170, well within the noise ceiling. As a disclaimer to our correlation analysis, we want to note that correlations computed on a small number of participants, e.g. less than 100, have typically low power [32,33]. However, here we are investigating within-subject correlations, which have typically higher power than between-subject correlations. Thus, the high correlation values and very similar effect topographies implicate the same face processing in passive and active contexts. This leads us to believe that the N170 and general lab-based face processing results generalize to more naturalistic setups, indicating that the found effect truly holds in everyday vision.

An important property of our study is that we investigated classic and naturalistic experimental settings within the same subjects. Besides the robust correlation of the N170 amplitude across the traditional and the naturalistic experimental conditions, we found that the between-subject variance in the naturalistic condition is smaller than in the classic setup. This came as a surprise, as the naturalistic setup contains more sources of variation, e.g. different gaze trajectories by different subjects. Thus, it appears that the difference in the visual processing of faces versus non-faces is more comparable across subjects under naturalistic conditions. As a note of caution, with the currently available data, we cannot make a definite statement with regards to the effect size. Future studies with more subjects might allow investigating the effect of ecological validity on inter-subject consistency. A highly speculative interpretation of this observation is that evolutionary constraints act under naturalistic conditions. To process the isolated image of a flashed face has no direct consequences for evolutionary success. However, the active fixation and visual processing of faces under naturalistic viewing conditions are arguably more directly related to relevant social interactions. That is, due to evolutionary constraints visual processing might be more consistent between humans under relevant naturalistic conditions as compared to artificial situations that the experimental subjects did not encounter before. If this speculative interpretation holds up in other studies as well, it would be a strong argument to investigate sensory processing under naturalistic conditions in general.

### Sequential effects during trajectories of fixation points

Our free viewing paradigm allows us a deeper insight into brain function, by analyzing sequential effects of fixation history. Our results show a positive shift in the ERP beginning before the current fixation for fixations originating from a face being more positive. This effect cannot be explained by a parafoveal preview [34], as it is only dependent on the category of the previous fixation, and not the current one, and no interaction between the two was found in that period of time. This effect might be attributed to neuronal fatigue. After processing a face during the previous fixation, the face processing system might exhibit a depletion. This depletion might override the negative effect of the source usually associated with the N170, which in turn will lead to a generally more positive activation. This amplitude reduction is in line with previous studies [24,34]. Notably, the fERP in our study is changing earlier than previously reported [35].

In the cases when the same face was fixated consecutively, we found an ERP difference to between-face fixations. This difference in activity could be due to adaptation effects to the specifics of the face, extending over and beyond the interaction of the previous and the current fixation previously described. Our finding contrasts those of Amihai et al. [36], who found no specific effect of identity repetition in a passive viewing paradigm. Interestingly, our finding extends to time points even before the onset of the fixation, potentially resulting from a type of within-face preview effect. Concerning this hypothesis, our results contrast those of previous studies which found no prefixation differences for congruent vs incongruent peripheral previews [34,37]. In our case, the participants were already looking at the face while refixating it, which might introduce an even stronger effect that is specific to free viewing paradigms. It is, therefore, a necessity to understand natural face processing in light of its recent history.

### A new methodology that allows for this type of analysis

In this study, we model both temporal overlap of neural processes in time, and non-linear influences of eye movement parameters (e.g. saccade amplitude or saccade position) which can lead to systematic differences between conditions [23,38,39]. Such regression-based deconvolution models are increasingly becoming popular (e.g. 23,40–43). The adequacy of our deconvolution approach can be seen in supplementary Fig.2, where we contrast it with a non-deconvolution analysis. Without overlap-correction, we see additional large differences for face vs. background already in the pre-fixation period and after 300ms. They can be attributed to two different overlap effects: The first effect is due to biased overlap with the stimulus response. The first fixations after stimulus onset are predominantly made on faces (nearly 70%, [44–46]). Thus, without overlap-correction, the fERPs of faces will be much more influenced by the stimulus ERP than background fixations leading to the observed bias. The second overlap effect is likely due to subsequent fixations. Fixation durations between faces and background fixation differed systematically, explaining this overlap effectv(see supplementary Fig.3). Besides previous simulation work, our results leave us confident that applying deconvolution and non-linear coefficient modeling is the right tool to analyze eye-movement-related potentials.

### Limitations of the present study

This study explores a new paradigm and naturally comes with limitations and unexplored questions. These questions pertain to the stimulus material used and eye-movements as quasi-experiments.

In our study, we used images taken from HD head-cams of freely moving participants. This has advantages and disadvantages. On one hand, the stimulus material is less well controlled. That is, the increased negativity of the N170 could be due to features like contrast, color, orientation, or luminance which might differ between faces and background fixations. While this is a point well taken, we have to recognize that faces do not come in e.g. all colors but are systematically different from the background, and faces are inherently more similar to each other than other objects [5]. Further, it should be noted that reduced control is an unpreventable result of studying vision in a more natural setting. Voluntary eye movements are a quasi-experimental setting that precludes randomization. Thus, causal statements like “fixating a face causes a larger N170” are more difficult to prove than in a classic experiment. Ultimately, we cannot completely exclude the possibility that an N170 evoked by eye movements is the result of a mediation effect induced by contrast or luminance, differences between fixation positions, as we cannot distinguish these factors with our dataset. On the other hand, the high diversity in our stimulus material in terms of low-level features has advantages as well. We present faces in a wide variety of viewing angles, distances, and, therefore, size, and lighting conditions. This natural variation leads to a lower within face similarity and thereby weakening the face similarity’s influence on the N170 amplitude. Further, it allows statements like “fixating a face under naturalistic conditions causes a larger N170 than fixating the background under these conditions.”

Additionally, we make use of a set of ecologically valid stimuli without photographer bias [45,47]. The consequence is that faces are viewed with many different sizes and from many different angles. This can be seen as a limitation, but we rather see it as a feature: if findings should generalize to other tasks and contexts, then they should be tested with a variable stimulus set. Because the typical cognitive neuroscience stimulus is quite specific, this lack of stimulus variability has recently been coined as the “generalizability crisis” [33]. Thus consequently, our statistical method should reflect the increased variability by addressing both between-subject and between-item effects. Unfortunately, it is currently computationally infeasible to adequately model this in combination with overlap correction as argued in [40], but see [48] for a counterexample. In addition, we argue that the very nature of natural stimulation necessarily implies a mixture of various signal sources, some of which can be artificial when experiments become more naturalistic and motion is allowed [49] but most of them are likely being used by the brain to extract meaning from the world.

Summing up, here, we advanced neuroscientific studies on face processing in multiple ways. We provide a naturalistic study setup, using natural scenes and allowing for eye movements, and combine this with an analysis pipeline that overcomes the technical challenges that are posed by this more natural setup. We reproduce previous findings from passive viewing and more controlled stimulus materials and show that the old and new effects closely relate to each other. Our findings also show that with a free viewing paradigm we can find previously unknown effects of eye movement history on ongoing face processing, opening new avenues of research for exploring vision in more natural, dynamic settings.

## Methods

### Participants

Twenty-three participants took part in our experiment. We excluded three subjects from further analyses. For one we could not synchronize the ET and EEG data. For the other two, the eye tracking data was not usable due to technical problems.

All 20 participants (15 female, 5 male; age: 19 to 31) reported normal or corrected to normal visual acuity. Participants gave written consent and were unaware of the purpose of the study. They received an hourly reward of either 8.84 € or course credits. The study was approved by the local ethics committee.

### Technical Setup

EEG data were recorded using a 128 Ag/AgCl-electrode system placed according to the 5% international system using a Waveguard cap (ANT, Netherlands) and two Refa8 (TMSi, Netherlands) amplifiers. We recorded with a sampling rate of 1024 Hz and used electrode Cz as the reference. The ground electrode was placed under the left collarbone. Eye movements were recorded via Electrooculogram (EOG) with a bipolar electrode being placed above and below the left eye. Impedances were kept below 10 kOhm.

Eye movements were recorded using an EyeLink 1000 remote eye tracker (EyeLink, SR Research, Canada) with a sampling frequency of 500 Hz in remote mode. For the passive viewing condition, we used a nine-point calibration before the first and fifth block. For the free viewing condition, we calibrated before the first, fourth, and seventh experimental block. The average calibration error was kept below 0.5° visual angle with a maximum error of 1.0°.

We used a large presentation screen with a width of 64” and a height of 36” (PA328Q, Asus, Taipei, Taiwan), a resolution of 3840×2160 pixels, and a refresh rate of 60 Hz. A luminance sensor was attached to the bottom left corner of the screen to detect changes in the monitor (i.e. stimulus on- and off-set). This was done to compensate for time delays between the trigger and the actual stimulus onsets. All data were corrected for this time delay. A jitter in this temporal delay was not found.

### Procedure

Participants were seated in a dimly lit room with their heads centered to the presentation screen at a distance of 80 cm.

The order of experiments was balanced between participants to avoid sequential task biases. Each experiment took about 40 to 60 minutes, including self-paced breaks after each block. The whole session including the EEG setup took about 3 to 4 hours.

#### Passive viewing

##### Stimuli

During the passive viewing condition, faces (front on or 40° rotation), objects, and cars were shown. For each stimulus category, we used twenty different identities. The face pictures were taken from a database of coworkers working at the NeuroBioPsychology Group of Osnabrück University. This database consists of photographs of 10 males and 10 females with neutral facial expressions, wearing black T-shirts with varying hair colors and styles from several different angles. Object photographs were taken from the Konkle’s Animacy x Size database [50] with 10 small and 10 large objects. Twenty car photographs were taken from [51]. These photographs depicted a range of different car types of various colors and shapes.

We matched the number of white pixels of each stimulus, but no other low-level features. Pictures could, therefore, vary in size. All stimuli were presented centrally on a bright white background. Car trials were part of a different research question and are not analyzed here.

##### Experimental design

Each passive viewing trial consisted of a fixation dot presented for 300ms followed by a stimulus presented for 300ms (Fig 4), followed by a white blank screen with an inter-trial-interval of 1300ms (uniform jitter of 1200-1400ms).

**Fig 4.**
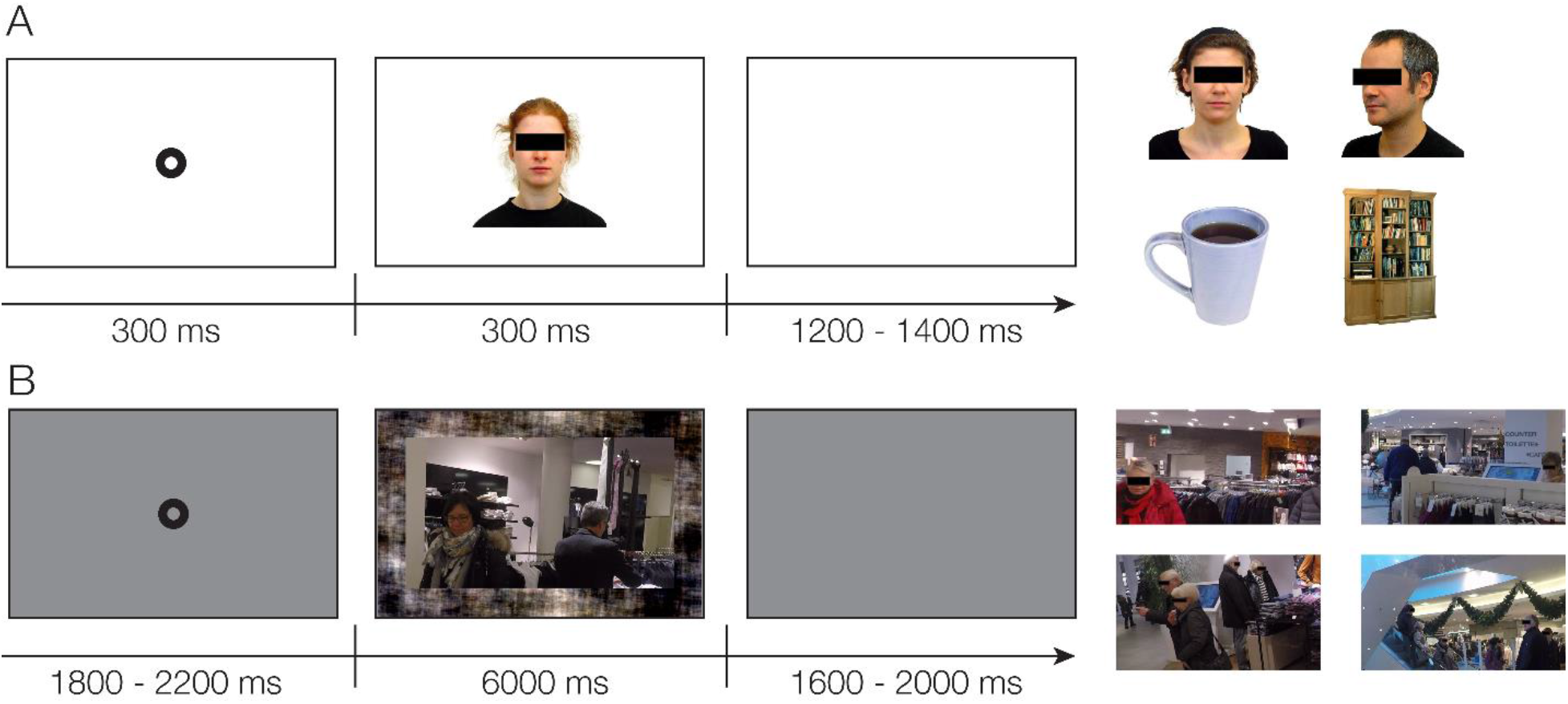
Exemplary Trial & Stimuli. (A) Trial structure of the classic, passive condition (left) and four exemplary stimuli (right). (B) Trial structure of the more natural, free viewing condition (left) and four exemplary stimuli (right).

**Fig 5.**
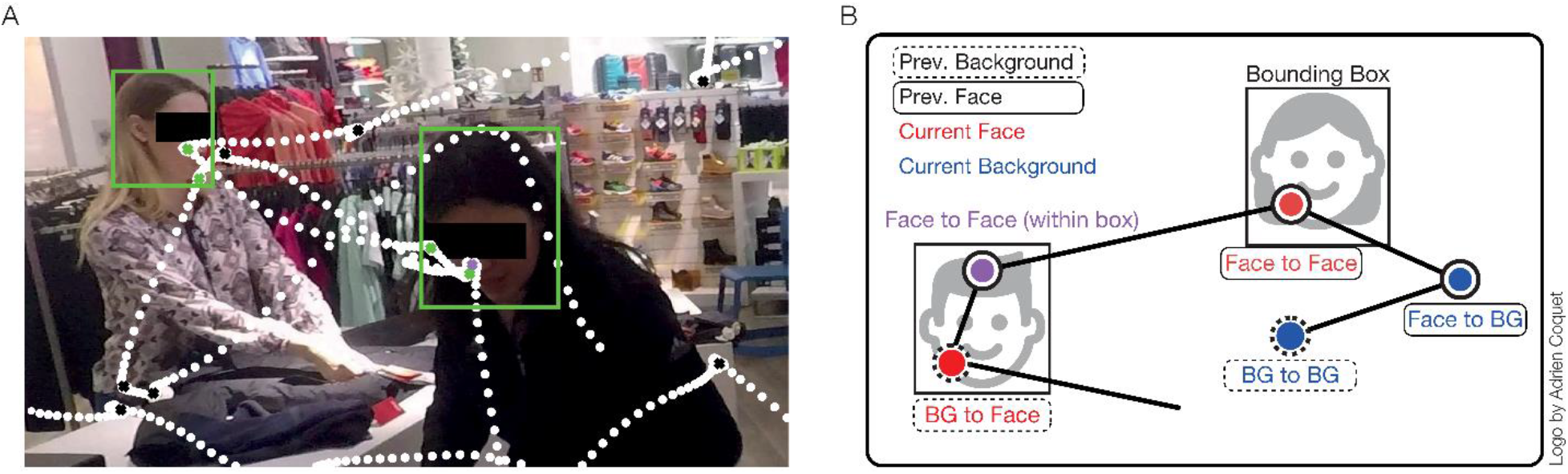
Exemplary eye tracking data of one trial and schematic visualization of the 2×2 categorization. (A) Eye tracking data of one subject. White dots represent the single samples, while the crosses represent the fixations as detected by the eye tracker. For visualization purposes, the faces are overlaid with their respective bounding boxes. (B) Fixations were categorized by their origin and their current placement. We distinguish between fixations made on the background (blue) or a face (red). For face to face fixations, we additionally specify whether they are the first fixation on a face, or a refixation within the same bounding box (within face fixations, purple).

We presented 1280 trials, where each of the 80 stimuli was presented 16 times randomly across eight blocks, except the half-profile stimuli, which were each repeated only 8 times per block. In total, there are 320 trials for each condition. The order of stimulus presentation within each block was pseudo-randomized, with no direct repetition of the same picture to avoid repetition effects. After every block, subjects were allowed to take a break.

#### Free viewing

##### Natural Stimuli

The stimulus set comprises scenes taken inside a local shopping center (Lengermann + Trieschmann, Osnabrück, Germany). To avoid the photographer’s bias, we recorded video streams with a GoPro camera (ASST1, Hero 5, GoPro, Inc., CA) mounted on a pilot subject’s head. The subject was freely moving inside the mall wearing a mobile EEG and ET setup and was given the task to explore.

We extracted single frames from the recorded video streams. In a first step, these frames were then manually screened and selected by criteria such as good visual acuity, straight camera angle, and the presence of faces. As subsequent frames are highly similar, in a second screening, we checked the images again and excluded similar-looking pictures to ensure a high stimulus-appearance variability. Next, we manually marked all faces in each frame with rectangular bounding boxes. We concurrently classified human faces, human heads (facing away), and non-human faces, like mannequins or faces on advertisements. In the experiment, all stimuli were displayed with a magnification factor of 2.53, in order to be perceived at the same size as in the real world.

Due to limitations of the eye-tracking device, accurate calibration could only be ensured in the inner 60% of the width and height of the screen (with the participant sitting 80 cm away). Therefore, the full screen images had to be cropped. To do so, we defined 25 overlapping sections placed in a 5×5 grid over each image. For each image, one of the sections was chosen as a stimulus by means of the highest number of human faces present. When more than one cut out contained the same number of faces, the section with the largest face was chosen. The stimuli contained between 1 and 7 human faces of different sizes and viewing angles. The size of the face annotation boxes ranged between 0.08° x 0.2° visual angle for the smallest and 5.2° x 5.6° visual angle for the largest box. This procedure resulted in two final sets of 171 images each. We presented each participant either the first or the second set, to minimize stimulus effects.

Ultimately, each stimulus was presented with a size of 30.5° x 17.2° visual angle. As they were not presented full screen, the remainder of the screen was filled with a phase-scrambled version of the respective image to minimize the effects of the fixation’s horizontal and vertical coordinate on the EEG signal [51].

##### Experimental Design

Each free viewing trial consisted of a fixation dot randomized between 1800 to 2200ms in the screen center, followed by 6000ms of stimulus presentation, and ended with a blank screen for a period randomized between 1600 and 2000ms. The experiment contained 9 blocks of 19 trials each, with self-paced breaks after each block. During stimulation, the subjects performed a free viewing task, being allowed to freely explore the presented scene. Subjects were previously informed that they are also allowed to look at the phase-scrambled background but that it did not contain any information.

At the end of each block, before the break, subjects performed a self-controlled guided viewing task. They would see 51 successive markers, randomly presented on a 7×7 grid, starting and ending with a marker in the screen center. Fixations of the respective marker were indicated by pressing the spacebar. These data are not analyzed here.

### Data Analysis

All analyses were done in MATLAB (Release 2016b, The MathWorks, Inc., Natick, Massachusetts, United States) using the EEGLAB toolbox v. 14.1.1b [53]. For integrating and synchronizing ET and EEG data the EYE-EEG toolbox (http://www2.hu-berlin.de/eyetracking-eeg) was used [38].

#### 2×2 statistical Design

Following the literature, we are interested in the difference between processing faces and other objects. In addition, we introduce sequential effects, as we hypothesized that the previous fixation category will influence the processing of the current fixation. This effectively results in a 2×2 design with the factors Current and Previous, both with levels Face and Object.

#### Eye Tracking

In both experiments, fixations were detected by the EyeLink system using the default cognitive setting (SR Research 2007). The eye tracker uses an acceleration-based algorithm to determine saccades, and fixations are classified as the non-saccadic segments. That is, fixations are defined by being below a certain threshold of acceleration within the eye tracker’s camera (velocity threshold: 30°/s, acceleration threshold: 8000°/s^2^, and motion threshold: 0.15°, [54]). Blink saccades, which were those spuriously detected due to blinks, were subsequently removed, by detecting whether a blink was enclosed between two saccades.

In the free viewing experiment, we identified the category of the currently fixated object and, to analyze sequential effects, of the previous fixation. We differentiated between i) fixations on a human face, ii) on a non-human face (mannequins, advertisements, etc.), iii) on a human head without a visible face, iv) on the background of the scene, or v) outside the stimulus on the phase scrambled border. Note that only fixations of type i) are of interest and all other types were not directly investigated here. Furthermore, we classified fixations whether they were on overlapping bounding boxes and whether consecutive fixations were within the same bounding box, i.e. within the same face.

While we estimated fERPs for all previously mentioned conditions, we focus on the previously introduced 2×2 design. In addition to the main effects of previous and current fixation-category and the interaction, we additionally investigated subsequent fixations on the same face.

#### EEG

##### Preprocessing

The eye tracking data was imported and synchronized with the EEG with the help of the EYE-EEG toolbox (v0.8) for EEGLAB [38].

Then EEG data were downsampled to 512 Hz and highpass filtered at 0.1 Hz (EEGlab plugin firfilt with a cutoff frequency 0f −6dB at 0.5 Hz, a hamming window, and a length of 3381 points, [55]).

Continuous data were visually inspected and artifactual sections were manually marked (muscle artifacts) and noisy channels removed (mean: 25.8, range: 19-34). Next, we used an independent component analysis (ICA, amica12, [56]) to remove components with eye-muscle artifacts [57]. Only for this step, the data were highpass filtered at 2 Hz to increase decomposition quality [58]. The ICA weights were then re-applied on the downsampled and continuous data. The ICA components were visually inspected and muscle and eye movement components were removed from the continuous data causally filtered at 1 Hz based on their topography, spectrum, and activation over time (mean: 22.41, range: 6-39). Data were re-referenced to average reference and removed channels were interpolated using spherical interpolation.

Because we need to correct for overlapping activity and eye tracking parameters, we used a regression-based approach implemented in the unfold toolbox [22]. A linear model including the factors Previous and Current (each consisting of the levels Background, HumanFace, and Other), the factor Samebox (if multiple fixations were made within the same face), and an interaction term was defined for the fixation ERP (fERP). Furthermore, spline regression was used to model non-linear effects of horizontal and vertical fixation position and saccade amplitude on the EEG. Additionally, the stimulus onset driven ERP (sERP) was modeled to correct for the overlap between the stimulus onset and the first fixation. This time expansion and thus overlap correction was applied between −500ms and 1000ms relative to fixation onset.

The data were modeled with the following Wilkinson-Formulas in the unfold toolbox by

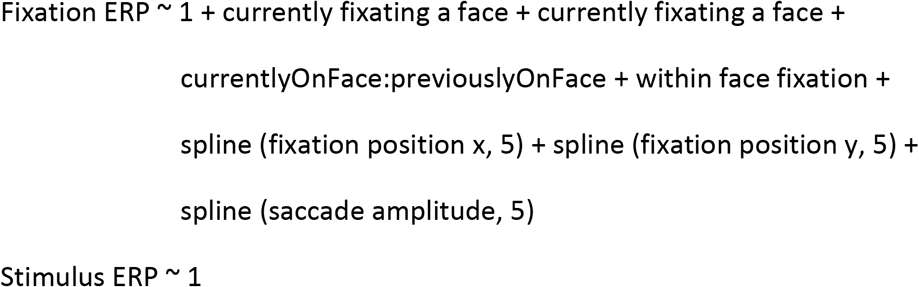

We used the same overlap correction for the passive viewing condition, even though we expected no overlapping activity between trials. However, participants did make some rare eye movements in the 300ms stimulus presentation, which might influence the ERP [23]; on the other hand, we keep comparability between conditions maximal by using the same analysis algorithms.

The passive viewing condition data were modeled with the following Wilkinson-Formulas:

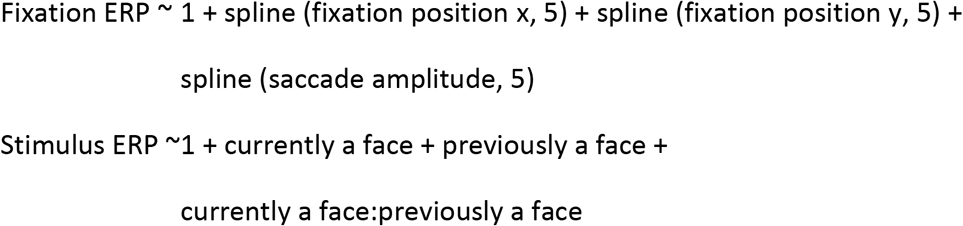

#### ERP Analysis

##### N170 analysis

The epoched, deconvolved ERP estimates were averaged over the occipital electrodes P7, PO7, P8, and PO8 according to [3]. The amplitude of the N170 was determined as the minimum in the time range of 130 to 200ms after fixation or stimulus onset according to [3], while the P100 was defined as the maximum between 80 to 130ms after the event of interest. After observing that some subjects had a P100 peak later than our initial prespecified time limit of 130ms, we extended the time limit to 150ms for all subjects. Additionally, in the lab condition, the N170 peaked earlier. Therefore, the time limits for the N170 were adjusted to 120 to 200ms.

##### Mass Univariate

Besides only performing the classic N170 analysis, we used the mass univariate approach to analyze the deconvolved ERPs for all electrodes and time points. Statistical testing was done using a one-sided t-test of parameter estimates at each time point with an alpha level of 0.05. The multiple comparison problem was corrected using a cluster-based permutation test with threshold-free cluster enhancement (TFCE) with 10.000 permutations. For each permutation, we randomly flipped the signs of each subject’s parameter estimate, calculated the t-values, and enhanced them using TFCE, generating an empirical H0 distribution of TFCE enhanced t-values. The maximum over the time range of −500ms to 1000ms was used to construct an H0 TFCE-value distribution, against which the actual TFCE enhanced t-values were compared. We considered t-values above the 95th percentile of this distribution to be significant.

##### Correlation and effect size

The correlation between the N170 amplitude from the passive viewing and Free viewing was calculated using the skipped Pearson correlation implemented in the robust correlation toolbox [59]. To minimize the effect of signal differences in previous time points, the peak-to-peak amplitude between the P100 and N170 was calculated [60]. We then subtracted the object peak-to-peak amplitude from the face peak-to-peak amplitude resulting in a difference-value for face processing for each subject in both the passive viewing and the Free viewing conditions. Seeing our high correlation value, we were interested whether this correlation value is compatible with a perfect correlation and calculated the noise-ceiling of an assumed perfect correlation, given the between (STD over subjectwise means, passive viewing: 1.6, free viewing: 1.0) and the within-subject variability (mean of subjectwise standard errors, passive viewing: 0.73, free viewing: 0.55). To simulate the between-subject variability, we sampled 20 new values from a normal distribution and scaled them each once by the condition-wise between-subject variability. This led to 2×20 values with a correlation of 1 (i.e. perfect). Because we cannot perfectly measure these data points, we added the within-subject sampling variability: for each subject and condition separately, we drew a random number from a normal distribution, scaled it by the respective within-subject variabilities, and added it. We repeated the procedure 1000 times, with each repetition resulting in a 2×20 matrix. For these randomly sampled results, we calculated the Pearson-correlation coefficient. The resulting distribution of Pearson correlations can be used as a parametric estimate of the H0 distribution taking measurement error into account. The median of this distribution is 0.8, whereas our observed correlation value is 0.78.

In order to calculate whether the between-subject variance in the free viewing condition was lower than in the classic lab condition, we bootstraped the individual experiment’s standard deviations. This procedure was done 10.000 times with all subjects detected as not being outliers in the robust correlation (see Fig 1C). Furthermore, we calculated the bootstrapped 95% confidence interval (10.000 repetitions) for the difference between the Cohen’s d_z_ of Passive Viewing and Free Viewing to estimate the difference in effect size with d_z_= mean(Face peak-to-peak - Object peak-to-peak)/std(Face peak-to-peak-Object peak-to-peak).

## Supplementary materials

**Supp. Fig 1.**
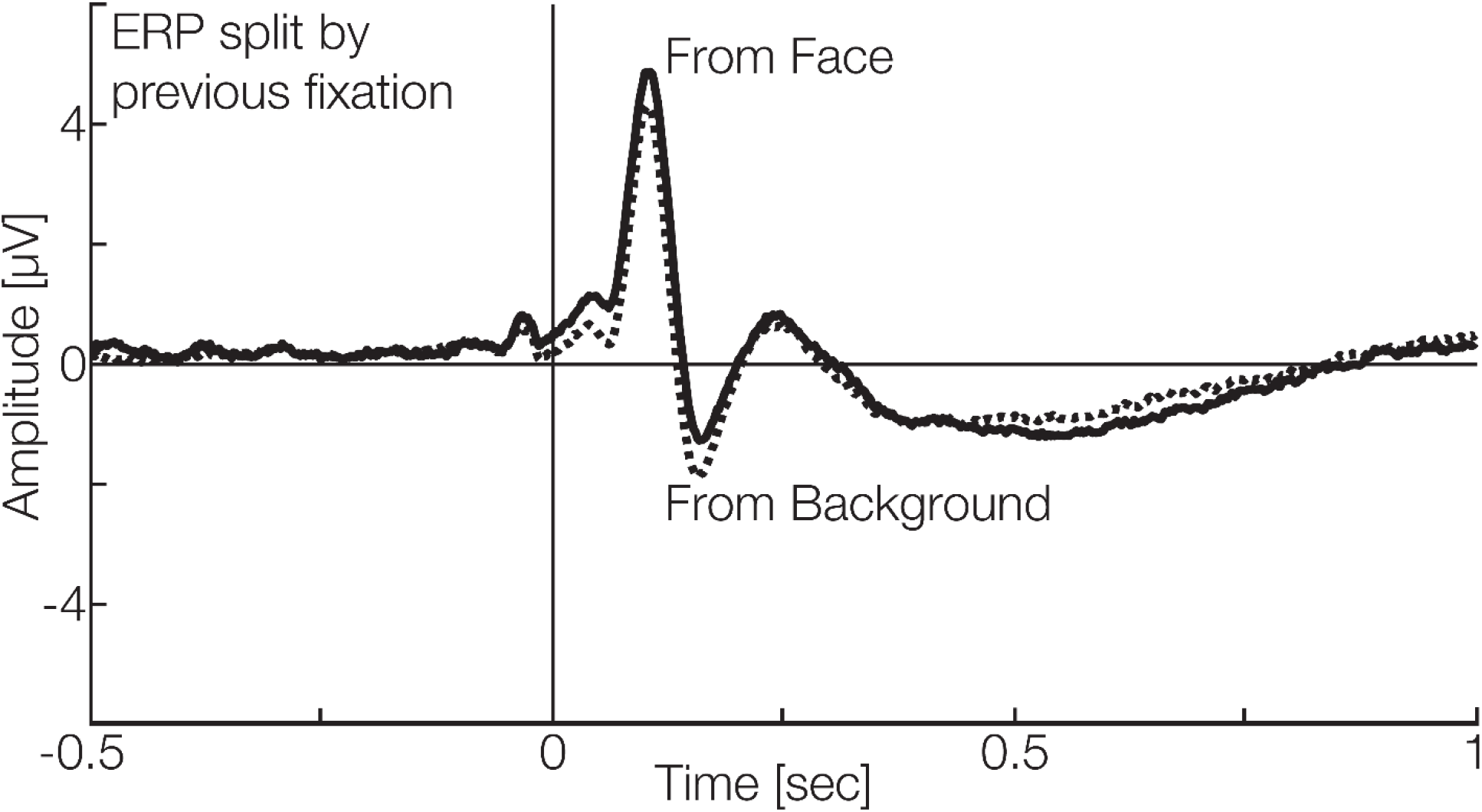
fERP split by the previous fixation. When coming from a face, the current fERP will show a positive offset independently of what is currently fixated.

**Supp. Fig 2.**
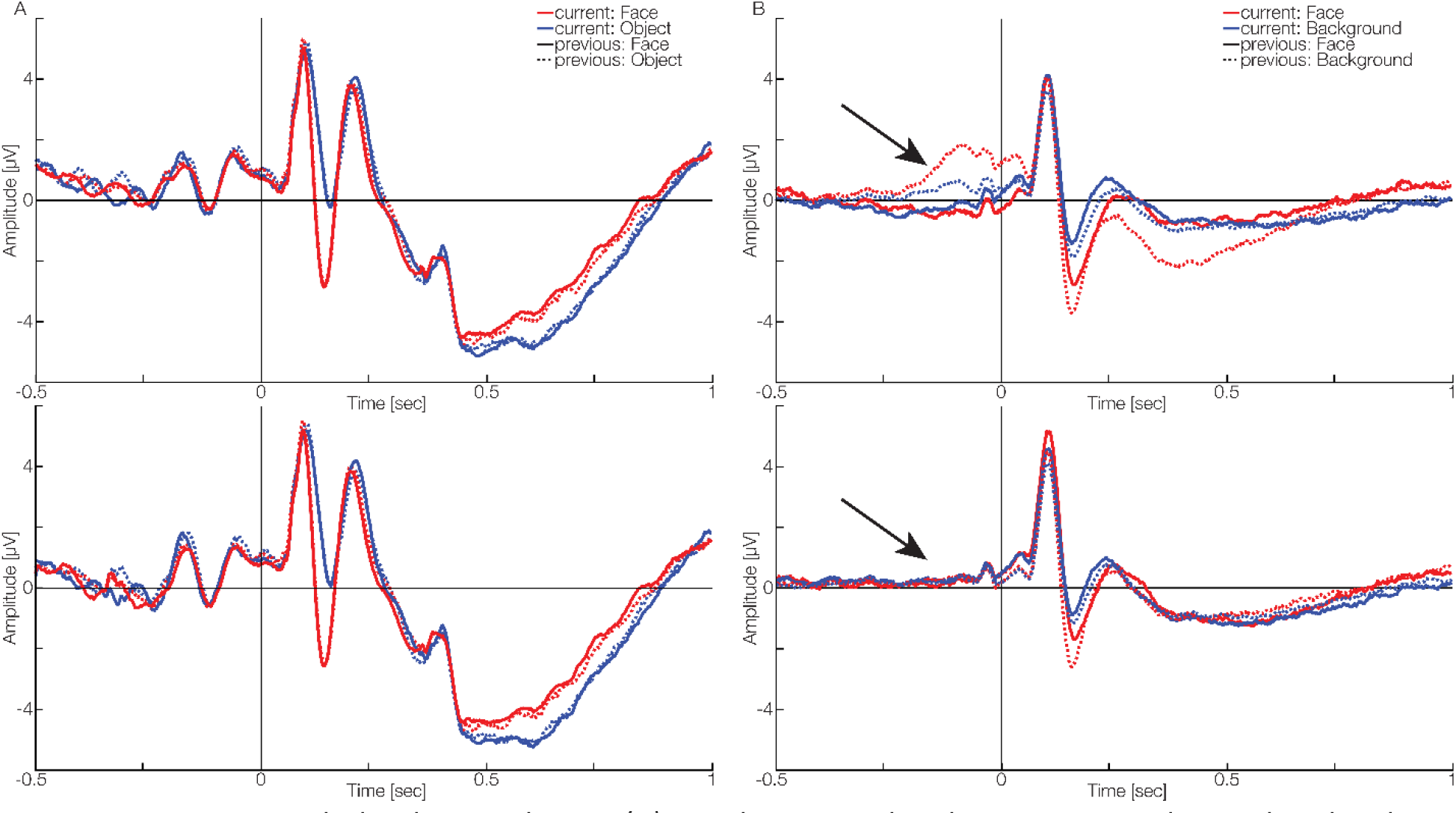
ERPs as split by the 2×2 design. (A) Resulting stimulus-driven ERPs as obtained in the classic lab condition. Top: Before the deconvolution, bottom: after. No major differences can be seen. (B) Resulting fixation-driven ERPs as obtained in the free viewing condition. Top: Before the deconvolution, bottom: after. Strong differences can be seen before the fixation onset. These spurious effects stem from overlapping activity of the stimulus onset. These changes can be explained by the dependencies on the eye movement parameters.

**Supp. Fig 3.**
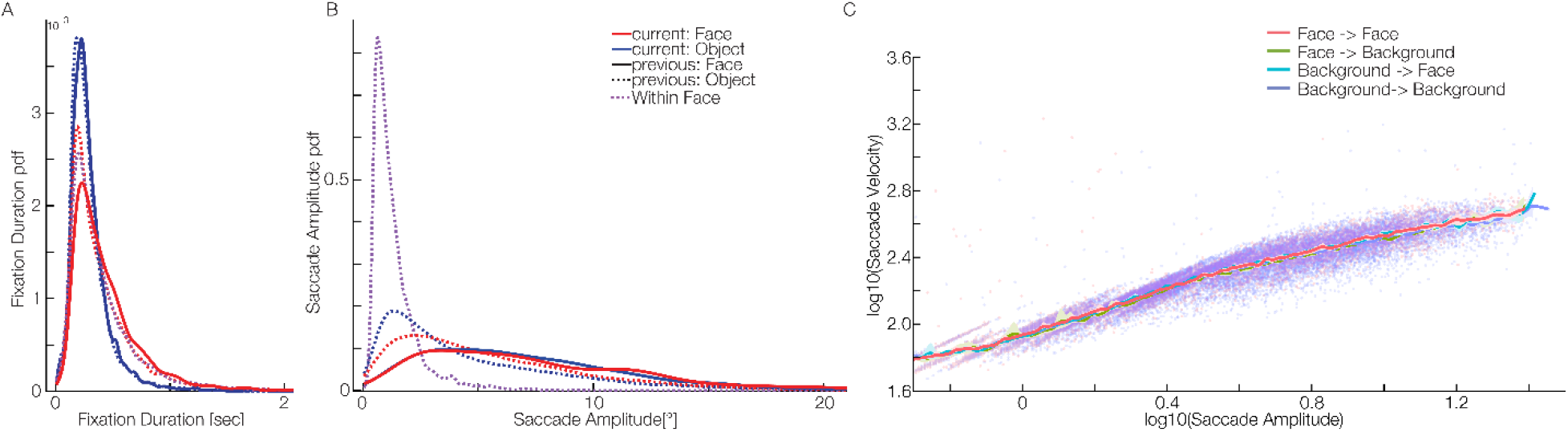
Distribution of eye movement parameters in the free viewing task as split by the 2×2 design. (A) Fixation duration. A clear distinction can be seen, leading to differences in overlap strength between the conditions. (B) Saccade Amplitude. Again, we see systematic differences, which might lead to differences in the ERP and therefore have to be controlled. (C) Main sequence. The eye tracking data show the typical main sequence.

**Table 1:** Details on the ERP amplitudes split by condition. Amplitudes and times are based on the individual participant’s ERP peaks. The confidence values are bootstrapped 10.000 times.

|  | P100 with 95%-CI | N170 with 95%-CI |
| --- | --- | --- |
| Obj to Obj | 6.4μV 95CI [5.0;7.7] at 104ms [98;108] | -0.97μV 95CI [-2.2;0.1] at 160ms [152;167] |
| Obj to HF | 6.2μV 95CI [5.0;7.5] at 96ms [93;100] | -3.6μV 95CI [-4.7;-2.4] at 143ms [138;148] |
| HF to Obj | 6.1μV 95CI [4.8;7.5] at 103ms [97;108] | -0.8μV 95CI [-2.1;0.3] at 156ms [151;161]. |
| HF to HF | 5.75μV 95CI [4.6;7.1] at 96ms [93;100] | -3.5μV 95CI [-4.5;-2.4] at 141ms [137;146]. |

Free viewing:
|  | P100 with 95%-CI | N170 with 95%-CI |
| --- | --- | --- |
| BG to BG | 4.5μV 95CI [3.7;5.5] at 102ms [97;106] | -1.6μV 95CI [-1.9;-1.3] at 162ms [154;170] |
| BG to HF | 4.9μV 95CI [4.0;6.0] at 101ms [97;105] | -3.0μV 95CI [-3.5;-2.6] at 155ms [152;161]. |
| HF to BG | 5.1μV 95CI [4.1;6.1] at 101ms [96;105] | -1.3μV 95CI [-1.7;-0.7] at 159ms [153;167] |
| HF to HF | 5.7μV 95CI [4.7;6.9] at 104ms [100;109] | -2.3μV 95CI [-2.7;-1.9] at 164ms [158;171] |

